# Composite Likelihood Method for Inferring Local Pedigrees

**DOI:** 10.1101/106492

**Authors:** Amy Ko, Rasmus Nielsen

**Affiliations:** Department of Integrative Biology, University of California, Berkeley, Berkeley, CA, USA; Department of Statistics, University of California, Berkeley, Berkeley, CA, USA; Museum of Natural History, University of Copenhagen, Copenhagen, Denmark

## Abstract

Pedigrees contain information about the genealogical relationships among individuals and are of fundamental importance in many areas of genetic studies. However, pedigrees are often unknown and must be inferred from genetic data. Despite the importance of pedigree inference, existing methods are limited to inferring only close relationships or analyzing a small number of individuals or loci. We present a simulated annealing method for estimating pedigrees in large samples of otherwise seemingly unrelated individuals using genome-wide SNP data. The method supports complex pedigree structures such as polygamous families, multi-generational families, and pedigrees in which many of the member individuals are missing. Computational speed is greatly enhanced by the use of a composite likelihood function which approximates the full likelihood. We validate our method on simulated data and show that it can infer distant relatives more accurately than existing methods. Furthermore, we illustrate the utility of the method on a sample of Greenlandic Inuit.

**Author Summary:** Pedigrees contain information about the genealogical relationships among individuals. This information can be used in many areas of genetic studies such as disease association studies, conservation efforts, and learning about the demographic history and social structure of a population. Despite their importance, pedigrees are often unknown and must be estimated from genetic information. However, pedigree inference remains a difficult problem due to the high cost of likelihood computation and the enormous number of possible pedigrees we must consider. These difficulties limit existing methods in their ability to infer pedigrees when the sample size or the number of markers is large, or when the sample contains only distant relatives. In this report, we present a method that circumvents these computational barriers in order to infer pedigrees of complex structure for a large number of individuals. From our simulation studies, we found that our method can infer distant relatives much more accurately than existing methods. Our ability to infer pedigrees with a greater accuracy opens up possibilities for developing or improving pedigree-based methods in many areas research such as linkage analysis, demographic inference, association studies, and conservation.

## Introduction

Pedigree information is used in many areas of human genetic analysis, including discovery of trait-associated markers in linkage analyses and family-based association studies [1], pedigree-informed phasing and imputation [2], and in estimating heritabilities [3]. Furthermore, pedigree information can be used to improve the performance of methods that otherwise assume unrelated individuals. For instance, in large association studies samples can harbor cryptic relatedness, which may result in spurious associations [4]. In such cases, pedigree information can be used to remove related samples or explicitly model relatedness to increase the power of association studies [5].

At the population level, pedigrees can elucidate the social organization and behavior of a group, such as mating patterns and variance in reproductive success among individuals [6]. Furthermore, pedigree information can be used to infer population parameters such as migration rates between subpopulations at very recent time scales. Most population genetic inference methods are based on coalescence theory, which models the genealogical relationships among samples of genetic data at a time scale of *N* generations. However, standard coalescence models, such as Kingman’s coalescent [7–9] ignore pedigree structure. Simulation studies have shown that the coalescent is a poor approximation of the genealogical process over short time frames (< log_2_N generations, where N is the population size), potentially leading to inaccurate inferences at these time scales [10, 11]. Therefore using the pedigree, which contains more detailed information about the genealogical history of the samples, should provide more power in inferring population parameters for the very recent past.

Pedigree information is undoubtedly valuable. In many cases, however, pedigrees are not directly observable and must be inferred from genetic data. Although numerous pedigree inference methods have been developed to date, most are limited to inferring very close relationships or require a prior knowledge of the sample structure. Many existing methods support only single-or two-generation samples. The single-generation methods are sibship inference algorithms which partition the sampled individuals into sibship clusters [12–15]. The parentage inference methods for two generations find the best parent-offspring combinations from a set of offspring and candidate parents [16–18]. Several methods that can support more than two generations have been developed [19–25]. But they are either limited in the number of markers that can be analyzed [20, 25]; do not support polygamous pedigrees [23, 24]; assume a complete sample (i.e. every member in the pedigree is sampled) [21, 22, 26]; or assume all sampled individuals belong to a single generation [23, 24]. The state-of-the-art method, PRIMUS [27], is the most flexible of the existing methods; it accommodates missing data and is able to infer multi-generational, polygamous pedigrees. Although PRIMUS is a notable improvement from other methods, its accuracy decreases significantly as the number of missing individuals increases. This is problematic as we expect samples to contain only a small fraction of pedigree members unless the sample represents a large portion of the total population or is specifically designed to include close family members.

The difficulty in pedigree inference comes from three sources. First, the number of possible pedigrees is enormous even for a small sample size [28, 29], making naive enumeration of pedigrees in search for the best one infeasible. Second, computing the likelihood of a pedigree is very expensive. Algorithms for computing the likelihood of a pedigree are either exponential in the number of loci [30], or in the number of individuals [31], which makes the likelihood computation of large pedigrees at many loci prohibitively slow. Finally, inference of pedigree relationships from genetic relationships, measured by the proportion of the genome shared by identical-by-descent (IBD), has high uncertainty. As the pedigree relationship between two individuals becomes more distant, the coefficient of variation and the magnitude of skew in genome sharing become larger [32]. For example, the distribution of genome sharing between second cousins overlaps significantly with that of third cousins, making these two pedigree relationships difficult to distinguish based on pairwise genome sharing alone.

In this report, we present a new pedigree inference method that addresses the drawbacks of the existing methods. More specifically, our method can utilize many markers genome-wide, support multi-generational pedigrees (up to 5 generations) and polygamous reproduction, and allows many missing individuals in the sample. We assume that all individuals are outbred and that the pedigrees do not create cycles, except in the case of full-sibs. To increase computation efficiency, we use a composite likelihood to approximate the full likelihood based on pairwise likelihoods, and use simulated annealing as a heuristic optimization algorithm for maximizing the composite likelihood. We validate our method on simulated data and show that it outperforms existing methods for inferring distant relatives. Furthermore, we demonstrate our method’s application to real data on a sample of Greenlandic Inuit.

## Materials and Methods

### Composite Likelihood

Our inference method is based on the idea of forming a composite likelihood function based on marginal likelihood functions calculated for pairs of individuals. While even pairwise likelihoods are slow to calculate for full genomic data, they can be tabulated and stored in computer memory. It is thereby possible to estimate pedigrees, based on a composite likelihood function, by only calculating the likelihood function between pairs of individuals once. This makes our method potentially applicable to large data sets containing thousands of individuals. As we will later discuss, using some heuristics, the method may even be applicable to large GWAS data sets.

We define a pedigree as undirected graphs where a node represents an individual and an edge represents a parent-offspring relationship (S1 Text). Each individual has a sex and is associated with 0, 1 or 2 edges connecting the individual to its parents, which mush be of different sexes if the individuals has two identified parents. An individual in the pedigree may or may not be represented in the sample, but if individual i is represented in the sample it is associated with genotype vector, *X_i_*. For each pedigree, the set of *k* sampled individuals is denoted by *H*, and the composite likelihood for such a pedigree is defined as

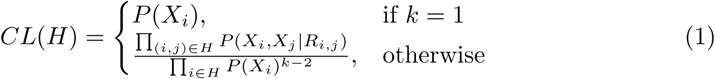

where *R_i,j_* is the relationship between *i* and *j* induced by the pedigree. For a singleton pedigree (i.e. pedigree consisting of one individual), the likelihood is simply the likelihood of the member individual’s genotypes. For *k* > 1 the composite likelihood is obtained as the product of marginal pairwise likelihoods. However, to obtain a more natural scaling of the composite likelihood we note that the probability of the data for each individual has been calculated *k* − 1 times and we therefore divide the composite likelihood function with the marginal likelihood of each individual *k* − 2 times. This has several desirable properties such as convergence of the composite likelihood to the true likelihood as the relatedness among individuals goes to zero. Another way to think of this composite likelihood function is in terms of products of conditional likelihoods. We can factor the full likelihood as

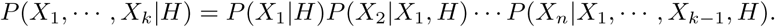

Since computing the conditional likelihoods *P (X_i_jX_1_, X_i1_; H)* is difficult, we approximate them with

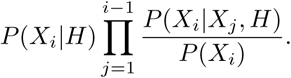

That is, we multiply the marginal probability of our current observation *P (X_i_|H)* by the likelihood ratio 
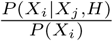
 for each previous observation *X_j_*. If the previous observation informs our current observation, then 
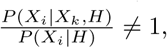

so the likelihood of the current observation increases or decreases accordingly. Using this approximation, we arrive at (1).

The pairwise likelihood *P (X_i_, X_j_|R_i,j_)* can be computed efficiently using the Hidden Markov Model (HMM) approximation by [33], which is used in this study. However, we note that any other definition of the pairwise likelihood function could have been used. For a set of possible outbred relationships in a 5-generation pedigree (See S1 Table), the pairwise likelihood for each pair (i; j) is precomputed and stored in memory. The total pre-computation time for 
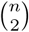
 pairs of individuals, *s* types of relationships, and *L* loci, therefore, is *O(n^2^sL)*. Since the composite likelihood of a pedigree is a simple function of the pairwise and marginal likelihoods, it can be computed fast by accessing the precomputed values stored in memory. The full composite likelihood for a set of local pedigrees is then computed by taking the product of the composite likelihood for each local pedigree.

It is worthwhile to note alternative ways to construct a composite likelihood. Another, perhaps more intuitive, formulation that also ensures that the composite likelihood converges to the true likelihood as the relatedness among individuals goes to zero, is

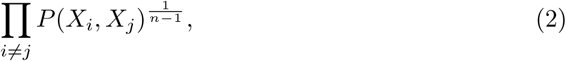

which scales the product of pairwise likelihoods by 
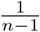
 to account for the multiple counting of each sample. However, as we will discuss in the Results section, this formulation leads to a worse approximation of the full likelihood function.

### Simulated Annealing

Because the number of possible pedigrees grows very rapidly with sample size, an exhaustive search for the most likely pedigree is infeasible for even a moderate number of individuals. Therefore, we use simulated annealing [34] to maximize the composite likelihood function. In this algorithm, a perturbation of the pedigree is generated by locally modifying the edges and nodes of the current pedigree (S1 Text). We explore the pedigrees with high likelihoods by always accepting proposals with higher likelihoods and occasionally accepting those with lower likelihoods to avoid getting stuck in local maxima. We implemented 24 different perturbations (moves) detailed in S1 Text. These moves can be broadly categorized into three classes. The first class of moves involves choosing two individuals and modifying their pairwise relationship. These moves include transitions between: parent-offspring and full siblings; full siblings and half siblings; first cousins and great avuncular; first cousins and half avuncular; and full avuncular and half siblings. Related to these are moves that add or subtract an edge between two nodes. For example, adding an edge causes parent-offspring relationships to become grandparent-grandchild relationship, whereas subtracting an edge has the opposite effect. The motivation for this class of moves is that these pairs of relationships have similar IBD coefficients, hence similar likelihoods. So these perturbations allow transitions between pedigrees with similar likelihoods.

The second class of moves allows bigger perturbations in the current pedigree. These moves include splitting a pedigree into two, joining two pedigrees into one, or the combination of splitting and joining. Splitting a pedigree can be done in two ways: we can either detach a chosen individual’s sub-pedigree (i.e. its descendant and itself) from its ancestors, or split off a randomly selected subset of its children to form a new pedigree. Joining two pedigrees involves creating a common ancestor between two individuals that belong to different local pedigrees.

The last class of moves is designed to transition between similar pedigrees when sex or age information is missing. For example, one move allows an individual and its descendant to swap places if age information is not present to resolve the directionality of the relationship. Another move changes the sex of an individual if sex information is not available, which in turn switches the sex of its potential spouses.

All of these transitions modify a small part of the current pedigree to generate a new configuration. Since the composite likelihood is a function of the pairwise and marginal likelihoods, the likelihood of the new configuration can be computed fast by adjusting the old likelihood by the changes made to the modified part of the pedigree.

The outline of the simulated algorithm is described below:

<u>Initialization:</u> Let each individual be a singleton pedigree (i.e. everyone is unrelated), except if there are known relationships that can be incorporated into the initial pedigree. Compute and store the composite likelihood of the current configuration.

<u>Recursion:</u>

1. Choose one of the 24 moves at random and generate a new configuration accordingly.
2. If the new configuration is an invalid pedigree, reject and go back to step 1. If it is a valid pedigree, compute the composite likelihood *CL(H_new_)* for the new configuration. Accept with probability *min[(CL(H_new_)=CL(H_old_))^t^*, 1], where *t* is the annealing temperature.
3. Repeat steps 1-2 C times.
4. Decrease the temperature to *t/f*, where *f* > 1 and go to step 1.

<u>Termination:</u> Terminate after *I* iterations or when the change in composite likelihood is less than e.

The tuning parameters *C, f, I*, and *e* were optimized for fast convergence using a number of trial runs on different simulated data sets. We run multiple instances of the algorithm with different random seeds. The algorithm then reports the pedigree with the highest likelihood encountered among all runs.

### Background Relatedness

Since the composite likelihood function is based on pairwise likelihood values, any inference based on it is limited by the quality of the pairwise likelihoods. One important factor that confounds the likelihood computation is linkage disequilibrium (LD), which often causes relationships to be overestimated [35]. Unrelated pairs of individuals often have higher likelihoods for being distantly related (S1 Fig), which leads to false detection of relatives. The method of [33] attempts to correct for LD by conditioning on nearby markers. However, in our experience residual effects of LD will still tend to bias inferences when markers are in high LD. One way to further reduce the effects of LD is pruning, or thinning, of markers. However, there is no consensus on how best to choose a set of markers that contains minimal LD and yet harbors enough information to detect distant relatives. To get a better sense of the effects of LD pruning on relationship inference, we simulated various pairwise relationships (i.e. second cousins, third cousins, unrelated) at linked loci. We then measured the pairwise prediction accuracy under different levels of LD pruning to choose an appropriate level of pruning threshold (See Results).

In addition to LD pruning, we further controlled for false detection of relatives by adding a regularization term to the composite likelihood. The regularizer was designed to weight against individuals from forming family clusters, motivated by the fact that in large data sets there are so many potential pedigree relationships for each individual, that most individuals will be inferred to have some pedigree relationship to at least one individual in the sample, even when they are unrelated. This is essentially a multiple testing problem in which an increasing number of individuals in the sample implies a reduced probability of inferring an individual to be unrelated to all individuals in the sample. There are natural ways of addressing this problem in a Bayesian framework that we might also be able to appeal to in the current framework. In particular, we will assign a probability distribution on the number of local pedigrees inferred. More specifically, we used the regularized composite likelihood

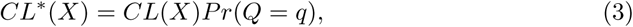

where *q* is the number of local pedigrees. We chose a Poisson distribution with mean *n*, the sample size, as the distribution of *Q*. This regularization is conservative in the sense that it favors every individual to remain a singleton unless there is strong evidence otherwise. Our choice to use the Poisson distribution was made, in part, for computational convenience but, as we will discuss in the Results section, resulted in good statistical properties of the method.

### Simulated Dataset

We tested the performance of our method on simulated pedigrees. We generated human autosomal haplotypes using msprime [36] with effective population size of 10,000, average recombination rate of 1.3e-8, and mutation rate of 1.25e-8. Using these founder haplotypes, we simulated three pedigree structures shown in Fig 1.

Simulation A consisted of 10 singletons and a 45-person family that spanned 5 generations. Of the 45 family members, 10 were sampled and 35 were missing. The kinship coefficients of the sampled relative pairs ranged from 1/4 (e.g. full siblings) to 1/256 (e.g. third cousins). Simulation B was designed to study the performance of our method on smaller family clusters. It consisted of 4 family clusters and 4 singletons. Each family cluster contained 15 to 18 members, of which only 4 of them were sampled. The sampled individuals spanned multiple generations and formed pairwise relationships with kinship coefficients ranging from 1/4 to 1/256. Finally, Simulation C was designed the test the method on pedigree structures in which every sampled individual, excluding singletons, has at least one close relative in the data. It consisted of 9 singletons and a 16-person pedigree that spanned 5 generations. The 16-person pedigree contained 7 missing individuals and 9 sampled individuals, where each sampled individual formed a parent-offspring relationship with at least one other sample.

**Fig 1.**
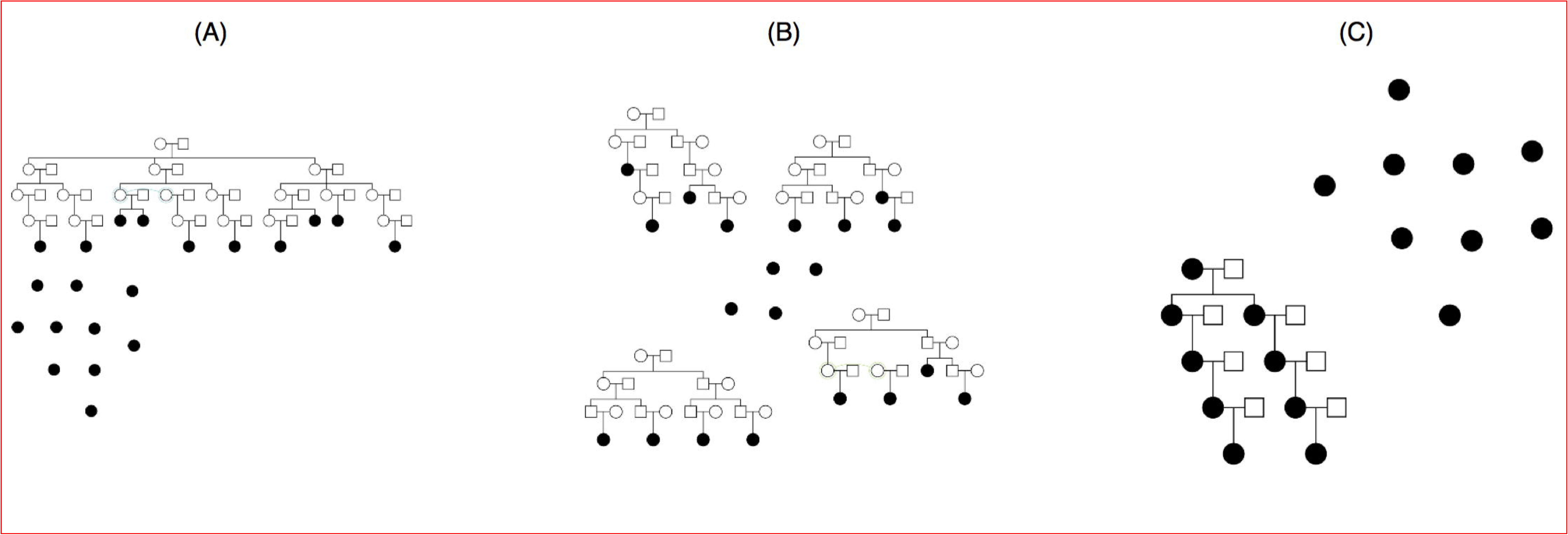
Simulated pedigrees. Shaded nodes indicate sampled individuals for which we have genotype data and unshaded nodes indicate unsampled individuals.

Each simulation scenario was replicated 100 times. For each sampled individual, we simulated genotyping error by switching each allele to the alternate allele with probability 0.01. To reduce the level of LD among markers, we used PLINK [37] to prune the original set of markers at *r*^2^ =:05, resulting in about 10,000 markers. The sex of each sample was assumed known, whereas the age was assumed unknown.

### Empirical Dataset

We applied our method to reconstruct the previously unreported pedigrees of 100 individuals in Tasiilaq villages in Greenland which had been genotyped [38] using the Illumina CardioMetaboChip, consisting of 196,224 SNPs. Since the European admixture into the Greenlandic population can confound relationship inference, we selected individuals from Tasiilaq villages, which showed one of the lowest levels of European admixture in the sample. In particular, the 100 individuals we selected were estimated to have European admixture proportion of 5 percent or less. To reduce the effects of LD, with pruned the markers using PLINK at *r*^2^ = 0:05. Due to the unusually high level of LD in the Greenlandic population, we were left with 1868 SNPs after LD-pruning.

### Competing Methods for Comparison

We compared the performance of our method on simulated data to PRIMUS, arguably the state-of-the-art pedigree reconstruction method. Although many pedigree inference methods exist, we chose to use PRIMUS as a benchmark since it is the most flexible of the existing methods in the types of pedigrees it can infer. More specifically, PRIMUS supports the inference of multi-generational, polygamous pedigrees and allows for missing individuals. PRIMUS reconstructs pedigrees that are consistent with pairwise IBD estimates and reports high-scoring configurations. To estimate the pairwise IBD coefficients for our simulated data, we first estimated the population allele frequencies from 200 simulated founder haplotypes. We then used PLINK to estimate the IBD coefficients for the individuals in our simulated pedigrees, where the population allele frequency estimates were provided as input. The IBD estimates were then used by PRIMUS to reconstruct likely pedigrees. This mimics the inference procedure recommended in the PRIMUS documentation.

We also compared our method to the pairwise inference method. In this method, we used the HMM by [33] to compute the pairwise likelihood under each possible relationship (S1 Table) for all pairs of individuals. Then we assigned each pair the relationship with the highest pairwise likelihood. We controlled the false positive rate by multiplying the likelihood of being unrelated by a scalar *c* > 0, in order to provide comparable results between methods. The pairwise inference method produces only the best relationship for each pair, which may not result in a valid pedigree when all pairwise relationships are pieced together. Still, it serves as a useful benchmark to evaluate the accuracy of pairwise predictions by our method.

### Measuring the Error Rate

We evaluated the performance of our method by comparing the pairwise relationships induced by the true pedigree to those induced by the estimated pedigree. We define the error rate for each pair as

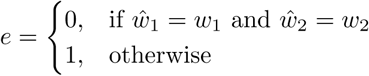

where *w_i_* is the probability that two individuals share *i* pairs of alleles IBD at a random locus under the true relationship; and 
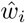

is the corresponding probability for the estimated relationship. In other words, the estimated relationship is correct if its three Jacquard coefficients [39] are exactly the same as those of the true relationship. For PRIMUS, which reports all pedigrees with the high likelihood scores, we compute the error rate by taking the average across all highest-scoring pedigrees.

Furthermore, to measure the distance between the estimated relationship and the true relationship for each pair, we compute the kinship coefficient distance

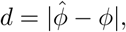

where 
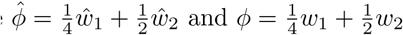

## Results

### Behavior of the Composite Likelihood

To examine the behavior of the composite likelihood, we simulated a nuclear family with two parents and their four children at 3,000 independent loci. We then computed the likelihood of the data under various pedigree configurations, ranging from the pedigree in which no one is related to the true pedigree. For each pedigree configuration, we computed the likelihood value with three different formulas: the full likelihood using MERLIN [40], composite likelihood A, given by (2), and composite likelihood B, given by (1).

The comparison of the three likelihood formulas are shown in S2 Fig. The x-axis is the distance of the test pedigree to the true pedigree, measured by the proportion of pairwise relationships that are correct in the test pedigree. As expected, the full likelihood increases as the test configuration becomes closer to the true pedigree. Both composite likelihood formulas preserve the ordering of the pedigrees induced by the full likelihood. That is, the order of pedigrees from the least likely to the most likely based on the full likelihood corresponds to the ordering based on the composite likelihood formulas. Although both composite likelihood formulas preserve this ordering, the likelihood surface given by (2) is much flatter than the full likelihood, whereas the likelihood surface of (1) is roughly on the same order of magnitude as the full likelihood.

### Effects of Linkage Disequilibrium on Pairwise Relationship Inference

As mentioned in the Methods section, we examined different thresholds for LD pruning. The appropriate level of pruning depends both on the genome length and the types of relationships we want to infer accurately. As shown in Fig 2, there is a trade-off between keeping enough markers to estimate distant relationships and removing markers to reduce false detection of relatives. For unrelated pairs, the most stringent LD pruning we tested (*r*^2^ =.025) showed the best relationship prediction accuracy. For third cousin relationships, however, pruning the markers too severely caused too much information loss, leading to a decrease in prediction accuracy. A similar pattern is observed for the second cousin relationships. For our simulated and empirical data, we prune the markers at *r*^2^ =0.05, which according to our simulations, retained enough information to estimate second and third cousins while keeping the false positive rate (i.e. estimating unrelated pairs as related) relatively low. We note that finding optimal strategies for dealing with background LD when inferring relatedness is an important topic that merits further research.

**Fig 2.**
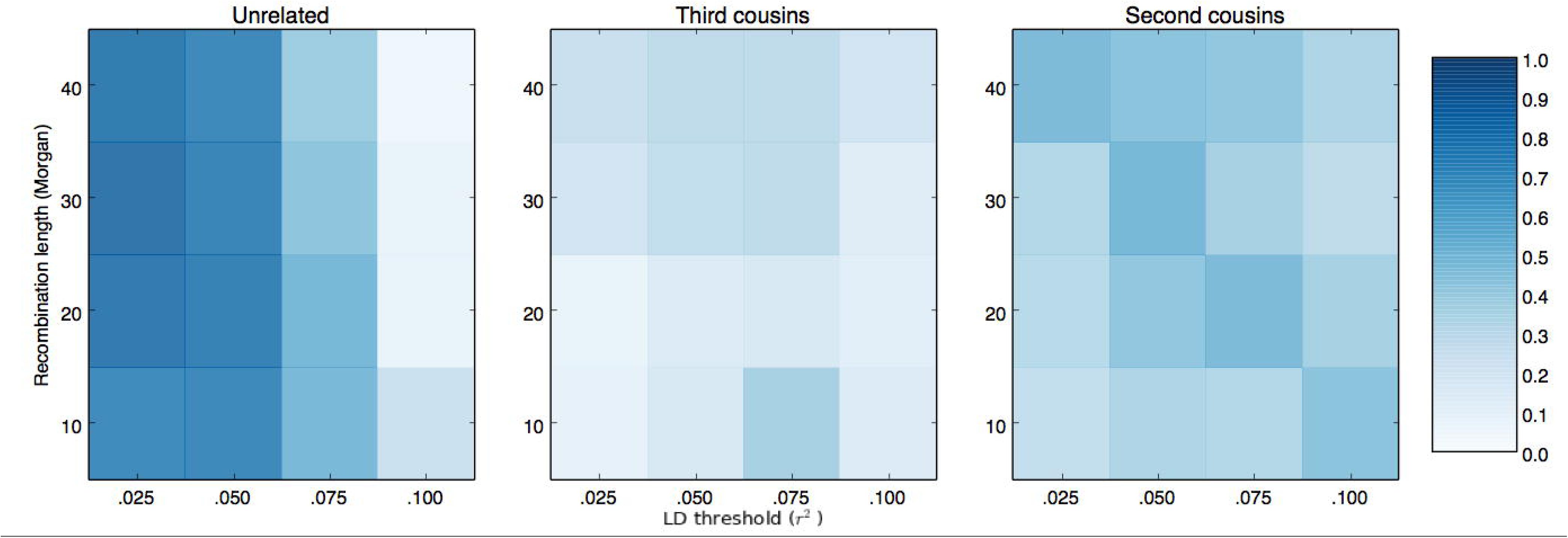
Effects of LD-pruning on pairwise prediction accuracy. The three panels show different true pairwise relationships: unrelated, third cousins, and second cousins. Each square in a panel corresponds to the relationship prediction accuracy for a particular genome length and LD-prune threshold.

### Estimating Simulated Pedigrees

Fig 3 shows the average pairwise error rate across all replicate experiments, categorized by different levels of true relatedness, *ϕ*. For Simulation A, both our method and PRIMUS had a very low false positive rate (i.e. error rate for *ϕ* = 0), and similarly low error rates for estimating close relationships. For more distant relatives such as first cousins and beyond ( *ϕ* ≤ 1/32), however, our method was able to estimate the relationships more accurately. PRIMUS estimated the distant relatives as unrelated in almost all instances, whereas our method was able to correctly infer such relationships about 50 percent of the time.

**Fig 3.**
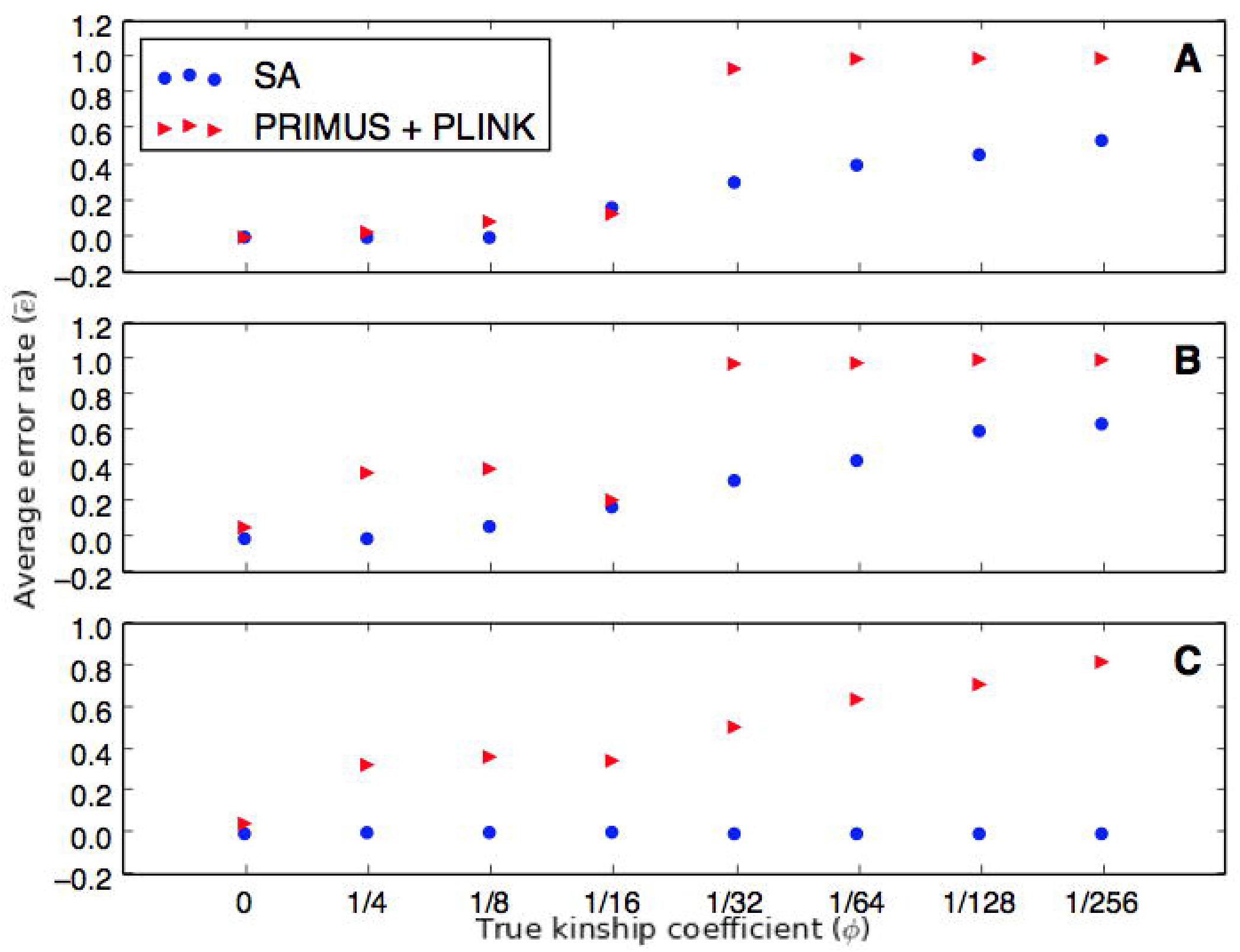
Comparison of prediction error rates between simulated annealing and PRIMUS. Each panel compares the average error rate between PRIMUS and our method (SA) for a particular simulation scenario: (A) Simulation A; (B) Simulation B;(C) Simulation C. In each panel, the x-axis shows different relationship categories measured by the kinship coefficient; the y-axis is the average error rate 
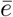
 (See Measuring the Error Rate in Materials and Methods section).

The middle panel in Fig 3 shows the error rate for Simulation B, which contains smaller family clusters than Simulation A. Again, our method outperformed PRIMUS in estimating relationships with *ϕ* ≤ 1/32. Unexpectedly, PRIMUS also showed higher error rates for close relationships such as avuncular relationship. This may be due to the uncertainty in pairwise IBD estimates, which PRIMUS uses to reconstruct the pedigree. With smaller family clusters, there are fewer pairwise relationships to inform and resolve more uncertain relationship assignments, which may lead to incorrect assignments of even close relationships. In that respect, simulation B is a more difficult pedigree to infer than Simulation A, and this is reflected in the error rates of our method as well, which are higher than the corresponding error rates in Simulation A.

For Simulation C, our method was able to find the correct pedigree in 96 of the 100 experiments. When the 4 experiments that converged to an incorrect pedigree were run again with a slower annealing rate, we were able to recover the correct pedigree in each case. The results for simulation C showed that when the sampled individuals are connected by close relationships, our method can unambiguously find the correct pedigree in most instances.

Our method also performed considerably better than the pairwise inference method. The likelihoods in the pairwise prediction were weighted so that its false positive rate roughly matched that of our method. Fig 4 shows that at similar false positive rates, our method estimated pairwise relationships with a greater accuracy than the pairwise method across almost all relationship categories. Fig 5 further demonstrates that our method has a significant advantage over the pairwise prediction method in detecting relatives. If the purpose of relationship inference is to find relatives-to discover the number of family clusters present in the data, for example-Fig 5 demonstrates that our method is able to detect relatives far more accurately than the pairwise method. These figures show that even though our method and the pairwise inference method both use the same pairwise likelihood values to estimate relationships, leveraging information from all pairs of relationships improves the inference significantly compared to considering each pair in isolation.

**Fig 4.**
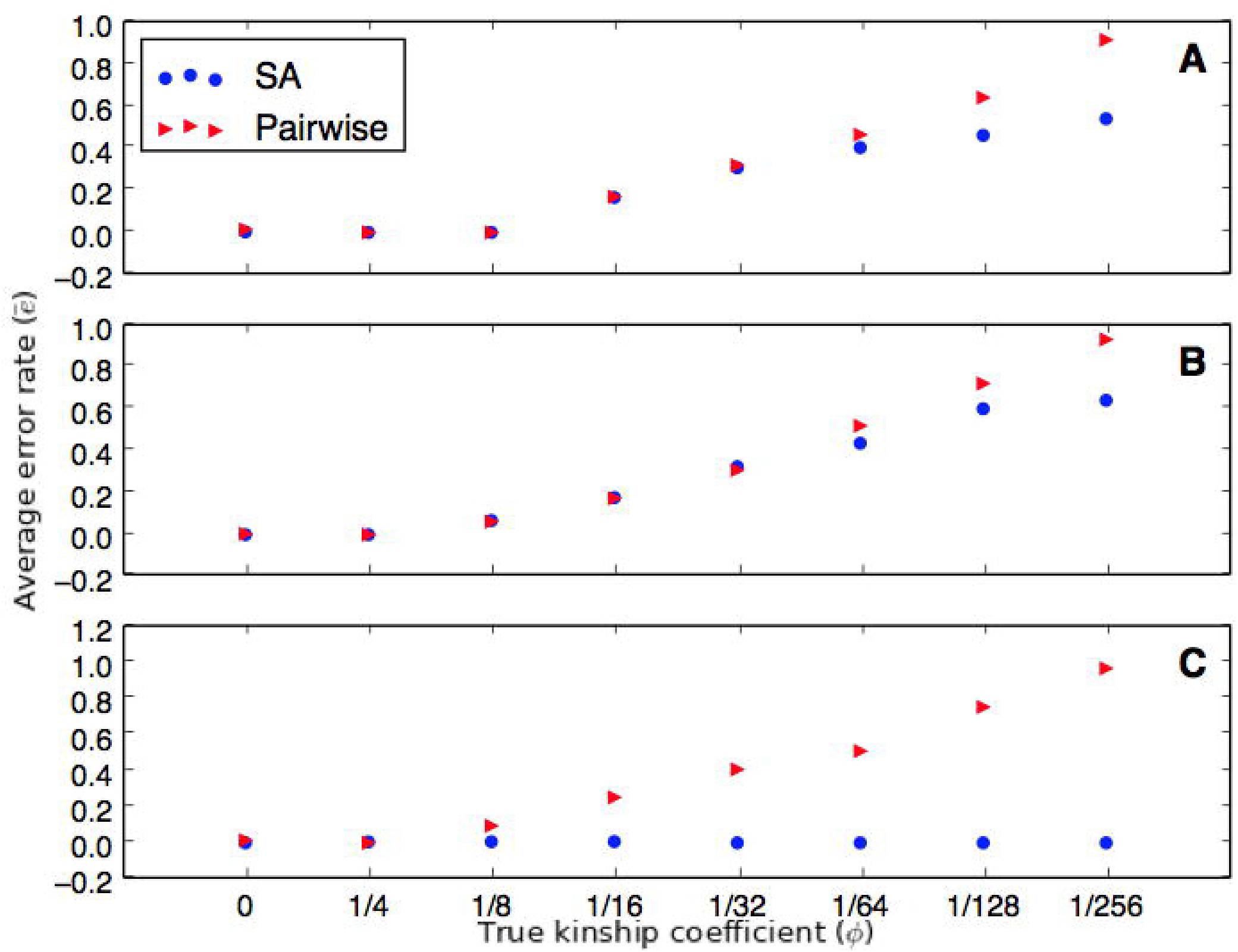
Comparison of error rates between simulated annealing and pairwise inference. Each panel compares the average error rate between the pairwise method and our method (SA) for a particular simulation scenario: (A) Simulation A; (B) Simulation B; (C) Simulation C.

**Fig 5.**
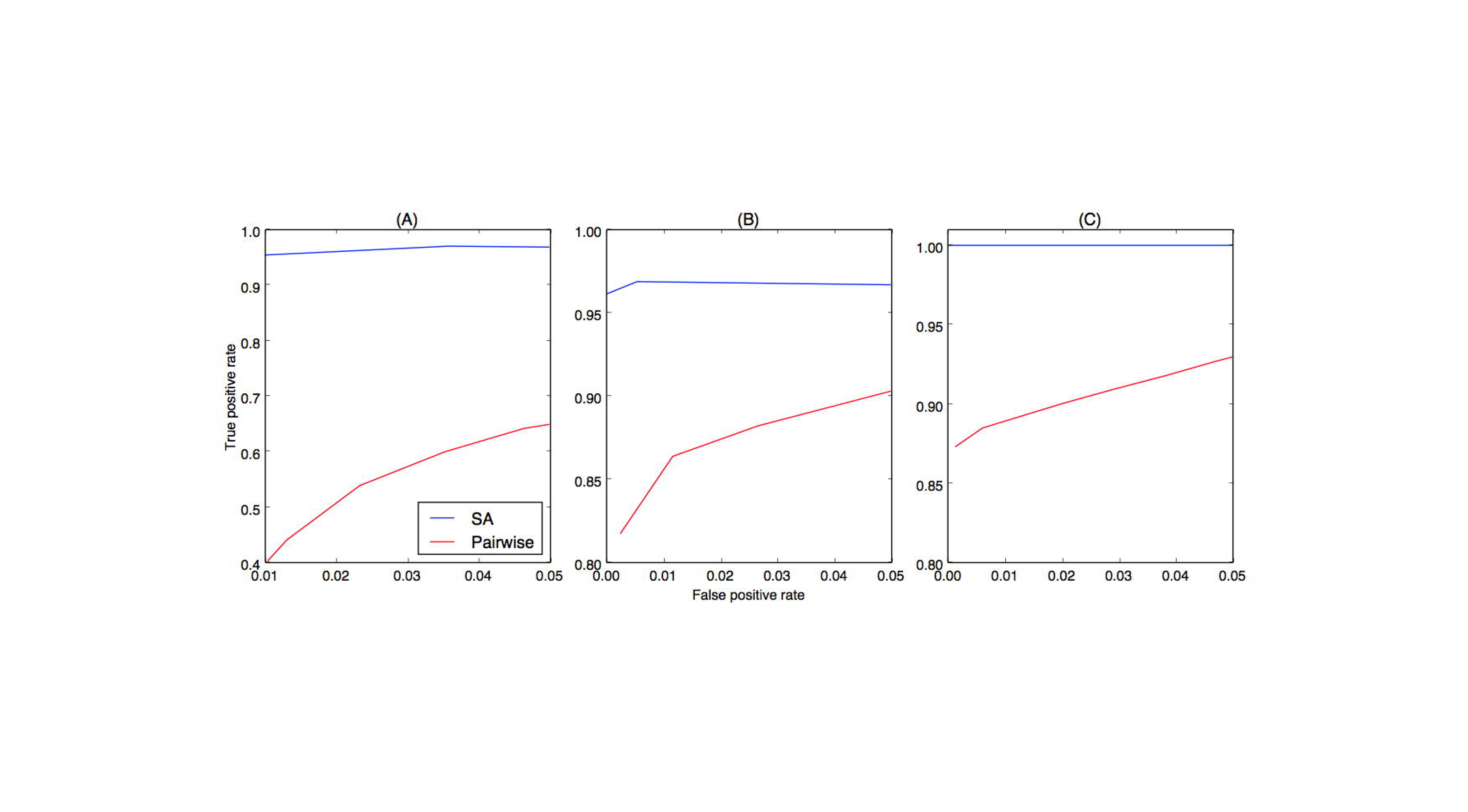
ROC curve for detecting relatives in a sample: pairwise vs. our method (SA). (A) Simulation A; (B) Simulation B; (C) Simulation C.

Furthermore, Fig 6. shows that even when the estimated relationship is wrong, it is generally close to the true relationship. For example, the maximum distance encountered in category *ϕ* = 1/32 was about 0.03 which is equivalent to predicting a first cousins once-removed relationship as first cousins. For the most distant relationship category we considered, *ϕ* = 1/256, most errors came from estimating these relationships as unrelated. The outliers in Simulation C resulted from 4 percent of experiments that converged to an incorrect pedigree in which a parent-offspring relationship was estimated as avuncular, thereby propagating errors down other related individuals.

**Fig 6.**
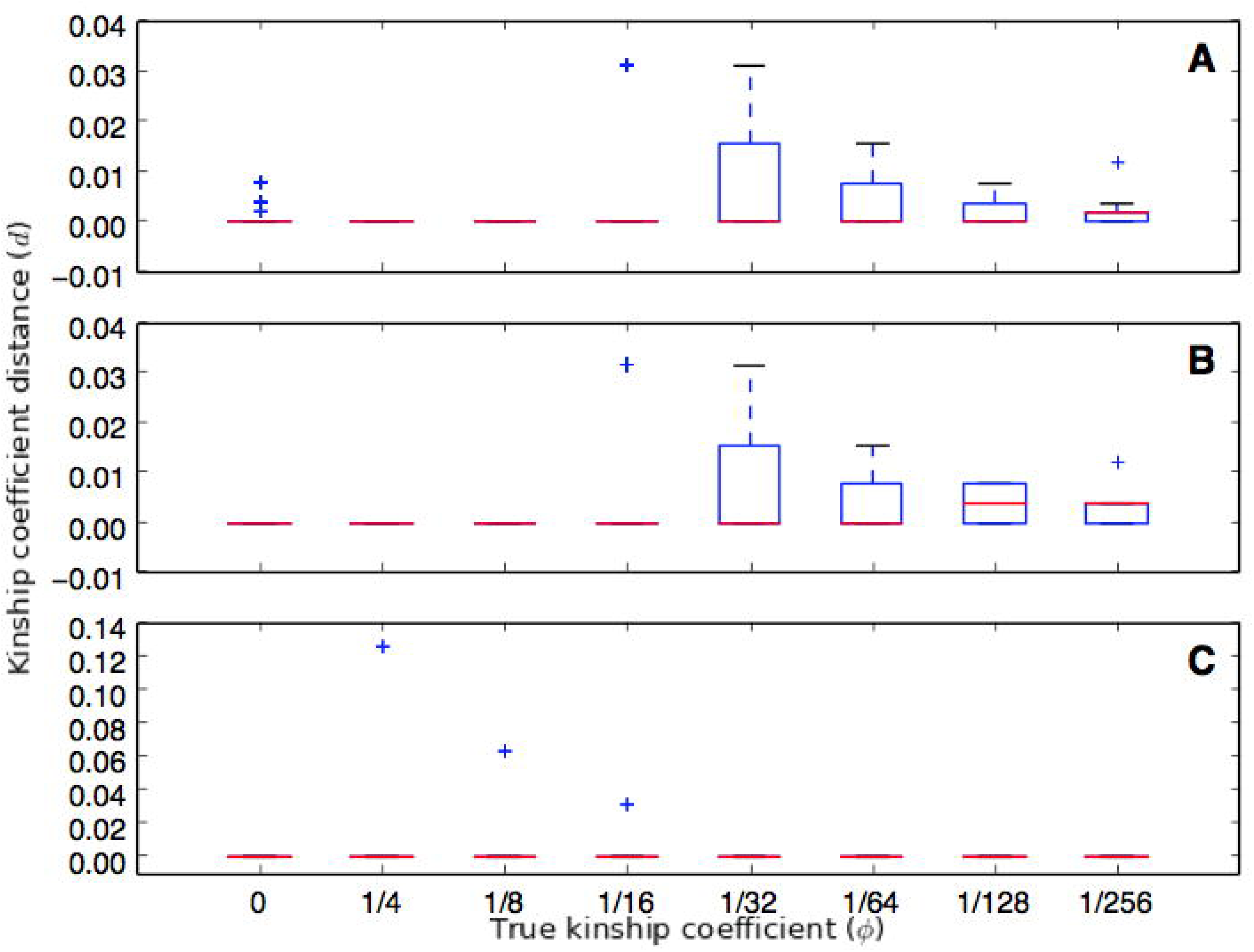
Absolute distance between the expected kinship coefficient under true and inferred relationships. (A) Simulation A; (B) Simulation B; (C) Simulation C. The x-axis is the relationship category measured by the kinship coefficient; the y-axis is the distance *d* between the true relationship and the relationship estimated by our method (See Measuring the Error Rate in Materials and Methods section).

Each experiment was run 3 times with different random number seeds, where each run consisted of 80 million iterations. The runtime of our method depends on many factors, including the number of individuals, the hidden pedigree structure, the number of missing individuals, and the annealing schedule in the simulated annealing algorithm. That said, each run on our simulated data took about 90 seconds on 2.5 GHz Intel Core i5 processor.

### Estimating the Greelandic Inuit Pedigrees

To demonstrate our method’s ability to infer pedigrees in practical applications, we estimated the previously unreportesd pedigrees of 100 individuals from Tasiilaq villages in Greenland. Because the Greenlandic Inuit population has high levels of LD, only 1868 SNPs remained after pruning the markers at *r*^2^ =:05. Our simulation study showed that at this number of SNPs, regularization with *Poi(n)* caused the error rate for estimating distant relatives (Φ = 1/32) to be very high; but using no regularization at all led to a high false positive rate (S3 Fig). So we chose to use *Poi(n/2)* as our regularization, which still produced a lower false positive rate, yet performed better in inferring distant relatives on simulated data.

We ran our algorithm 5 times with different random number seeds. Each run, which consisted of 80 million iterations, finished in about 24 minutes on 2.5 GHz Intel Core i5 processor. We used CraneFoot [41] to draw the estimated pedigree (S4 Fig). The reconstructed pedigree consisted of 38 singletons and 8 non-singleton family clusters. Many of these clusters consisted of close relationships such as parent-offspring, full siblings, half-siblings, and avuncular relationships. Based on our simulations, we expect more than 90 percent of the estimated relationships in these categories to be correct.

## Discussion

Our method provides a computationally tractable way to estimate pedigrees for a large number of individuals at many loci. The use of composite likelihood allows us to analyze pedigrees containing many individuals at many loci, where computing the full likelihood would be prohibitively slow. Furthermore, our method can estimate pedigrees when the number of possible pedigrees is too large to enumerate, which is true even for tens of individuals in a multi-generational pedigree. Our method is also one of the very few methods that can support complex pedigree structures such as polygamy, multigenerational pedigrees (up to 5 generations), and missing individuals. In addition, we can incorporate information about sex, age, and the number of generations spanned by the sample to better estimate the pedigree.

We have shown that our method has a significant advantage over the pairwise inference method. It can better estimate relationships beyond first cousins (Fig 4) and is able to detect relatives much more accurately (Fig 5). The composite likelihood considers all pairwise likelihoods jointly, which in turn can help resolve uncertain relationships in the context of other pairwise relationships. Therefore, even for pairwise relationship inference, where estimating the entire pedigrees may not necessarily be of interest, our method can be used to estimate the relationships more accurately.

Our method also showed an improvement over PRIMUS, the state-of-the-art pedigree reconstruction method, in inferring distant relatives when the number of missing individuals is large. PRIMUS’s reconstruction algorithm relies on accurate pairwise relationship assignments based on IBD estimates. If the sample consists mostly of distant relatives, however, relationship assignment becomes uncertain due to high variance in IBD sharing, which often leads to incorrect pedigree reconstruction. Although our method also relies on pairwise information, we showed that working directly with pairwise likelihood values rather than IBD-based relationships assignments improved the power significantly. Furthermore, PRIMUS’s enumeration of possible pedigrees becomes computationally cumbersome as the number of likely pedigrees increases rapidly for a set of distantly related samples. If the data contains many close relationships, however, PRIMUS can reconstruct all likely pedigrees very fast, whereas our method produces a single best pedigree, which may be close but not exactly correct. Thus the performance of each method depends on the sample structure and a suitable method must be chosen accordingly.

We applied our method on the Greenlandic Inuit dataset to demonstrate its ability infer previously unknown pedigrees from genetic data. Although the estimates of distant relationships are uncertain, we can still get a general sense of pedigree structures hidden in the data and take appropriate actions for downstream analyses. For example, the inferred pedigree can be used to filter out close relatives or model relatedness among samples in association studies. Furthermore, we can validate or improve the estimated pedigree with other evidence such as age.

Pedigree inference based on our composite likelihood is heavily influenced by how well we can compute the pairwise likelihoods. An important factor that affects the pairwise likelihood computation is LD, which often leads to overestimation of relatedness. Although the HMM by [33] conditions on nearby markers, it does not remove the effects of LD completely and necessitates LD-pruning. Unfortunately, there is no consensus on how best to prune markers while still retaining enough information to infer distant relatives. Although we carried out a simple simulation study to get a rough sense of appropriate level of pruning, it is by no means a complete solution. More work is needed on the effects of LD on relatedness inferences and how to remedy the problem, whether it be by more extensive simulations studies, or by modeling LD in the likelihood computation. Furthermore, care must be taken to use appropriate allele frequencies in likelihood computation to account for other potentially confounding factors such as population substructure [42, 43] and admixture [44, 45]. As better methods for estimating pairwise likelihoods become available, our method for estimating pedgirees should also improve.

There are limitations of our method that require further work. Our method assumes that all individuals are outbred, which may not be true of many systems including some human populations [46, 47]. It currently does not support pedigrees with cycles caused by inbreeding or complex cyclic relationships such as double first cousins. Another limitation of our method is that it does not provide any uncertainty measure on the estimated pedigree. A possible solution to this problem is to extend our method to a Monte Carlo Markov Chain (MCMC) algorithm to estimate the posterior distribution of the pedigrees. Casting our method in a Bayesian framework would also allow us to use a prior distribution to control the false detection of relatives. Furthermore, while computationally efficient compared to full likelihood methods, our method is still based on calculation of pairwise relationships and does, therefore, not scale up to GWAS data sets with hundreds of thousands of individuals. However, it may be possible to use a divide-and-conquer approach in which individuals are first divided into clusters using methods such as [48], then estimating the pedigree of each cluster separately, and finally estimating more distant relationships among clusters.

Overall, our method provides a computationally efficient way to estimate pedigrees of seemingly unrelated individuals. It improves our ability to validate and discover pedigrees in realistic genetic datasets where we expect a high level of missing data. The ability to estimate pedigrees more accurately opens up possibilities to develop and improve numerous pedigree-based or pedigree-aware studies, from correcting cryptic relatedness in GWAS to estimating demographic parameters of the very recent past.

Our software is available for download at https://github.com/amyko/pedigreeSA.

## Supporting Information

**S1 Table. Summary of Possible Pairwise Relationships in a 5-generation Pedigree.**

**S1 Text. Description of Transitions between Pedigree Graphs.**

**S1 Fig. Effects of LD on Relatedness Estimation.** The figure shows the histogram of the log likelihood difference, L(unrelated) L(third cousins), when the true relationship is unrelated. Unrelated pairs often have higher likelihoods for being third cousins when LD is present in the data, as shown by the histogram corresponding to linked markers. The data were simulated with msprime and the full likelihood was computed using MERLIN.

**S2 Fig. Comparison of Various Likelihood Formulas on Simulated Data.** The x-axis measures how close the test pedigree is to the true pedigree; the test pedigree becomes closer to the truth from left to right. In this simulation, the composite likelihood given by (1) approximates the full likelihood more closely than (2).

**S3 Fig. Effects of Regularization Term.** Accuracy of simulated annealing method on simulated data at 2000 markers under different levels of regularization.

**S4 Fig. Estimated pedigree of 100 Tasiilaq Individuals in the Greenlandic Inuit Dataset.** Shaded nodes indicate sampled individuals; unshaded for unsampled; squares for male; circles female; diamonds for unknown sex. Any two individuals connected by colored lines indicate they are the same individual.

**S5 Fig. Likelihood Convergence for the Greenlandic Inuit Pedigree Estimation.**

## Acknowledgments

We would like to thank Jeffrey Spence and Peter Wilton for helpful discussion and feedback, and Matteo Fumagalli for providing the Greenlandic dataset.

